# Multiple Protein Structure Alignment at Scale with FoldMason

**DOI:** 10.1101/2024.08.01.606130

**Authors:** Cameron L.M. Gilchrist, Milot Mirdita, Martin Steinegger

## Abstract

Protein structure is conserved beyond sequence, making multiple structural alignment (MSTA) essential for analyzing distantly related proteins. Computational prediction methods have vastly extended our repository of available proteins structures, requiring fast and accurate MSTA methods. Here, we introduce FoldMason, a progressive MSTA method that leverages the structural alphabet from Foldseek, a pairwise structural aligner, for multiple alignment of hundreds of thousands of protein structures, exceeding alignment quality of state-of-the-art methods, while two orders of magnitudes faster than other MSTA methods. FoldMason computes confidence scores, offers interactive visualizations, and provides essential speed and accuracy for large-scale protein structure analysis in the era of accurate structure prediction. Using Flaviviridae glycoproteins, we demonstrate how FoldMason’s MSTAs support phylogenetic analysis below the twilight zone. FoldMason is free open-source software: foldmason.foldseek.com and webserver: search.foldseek.com/foldmason.

## Introduction

Multiple protein comparison is vital to elucidating the evolutionary history, diversity and function of proteins. Computing optimal multiple sequence alignments (MSA) has been shown to be NP-complete (1). Therefore, tools for multiple alignment adopt heuristics, such as progressive alignment (2). Here, a guide tree is first estimated from unaligned sequences, and then sequences are sequentially aligned following the order in the guide tree from most similar to dissimilar. Many progressive aligners have been developed to construct MSAs, notably MAFFT (3), MUSCLE (4), Clustal Omega (5) and FAMSA (6). The inclusion of structural information has been shown to produce higher quality alignments than with protein sequence alone (7–9). Protein structure is conserved beyond the level of amino acid sequence and allows for comparison of proteins that have diverged beyond the ‘twilight zone’ level of sequence similarity (10, 11). This has driven the development of several tools for constructing multiple structure alignments (MSTAs). Like sequence aligners, most structure aligners adopt the progressive alignment heuristic, sequentially building MSTAs from pairwise structure alignments (PSAs).

For instance, US-align (12, 13) generates PSAs of all pairs of input structures using TM-align (14), then progressively aligns the PSAs following a UPGMA guide tree. Caretta-shape (15, 16) uses counts of overlapping Zernike backbone fragments to generate a guide tree and align structures through superposition and dynamic programming. Both US-align and Caretta-shape consider amino acid sequence similarity. MUS-TANG (17) progressively aligns structures based on residue-residue contacts and local structural topology. In contrast, Matt (18) uses an aligned fragment pair chaining algorithm to allow for structural flexibility rather than a progressive alignment. These MSTA methods excel in accuracy at the expense of high computational demands. Recent breakthroughs (19, 20) in structure prediction now necessitate orders of magnitude faster methods to scale to databases containing hundreds of millions of structures such as AlphaFoldDB (21) and the ESMAtlas (20). We recently reported Foldseek (22), which can query massive structure databases and identify pairwise similarities. Fold-seek uses a novel structural alphabet (3Di+AA) to represent tertiary interactions within protein structures as one dimensional sequences, making them amenable to fast string comparison algorithms. We further extended Foldseek to structural clustering and applied it to the AlphaFoldDB to generate the AFDB clusters (23). These efforts resulted in clusters of protein structures containing up to hundreds of thousands of highly-divergent members. Fully illuminating these clusters requires accurate and scalable MSTA computation.

To tackle this, we present FoldMason (**Fig. 1a**), a novel progressive aligner. Using benchmarks, we show that FoldMason produces MSTAs of comparable quality to gold-standard, while operating two to three orders of magnitudes faster and approaching the speeds of the fastest sequence aligners. It first encodes protein structures to be aligned using the 3Di+AA alphabet, performs all-vs.-all comparison via a striped accelerated ungapped alignment (24, 25) and generates a minimum spanning tree that iteratively guides MSTA construction through a parallelized progressive alignment. In the first iteration, PSAs are computed from the 3Di+AA sequence; subsequent iterations pairwise align structure profiles built from previously merged sets of structures. Optionally, FoldMason performs a linear time-clustering (26) as a preprocessing step to form preliminary groups and improve its scaling.

**Fig. 1.**
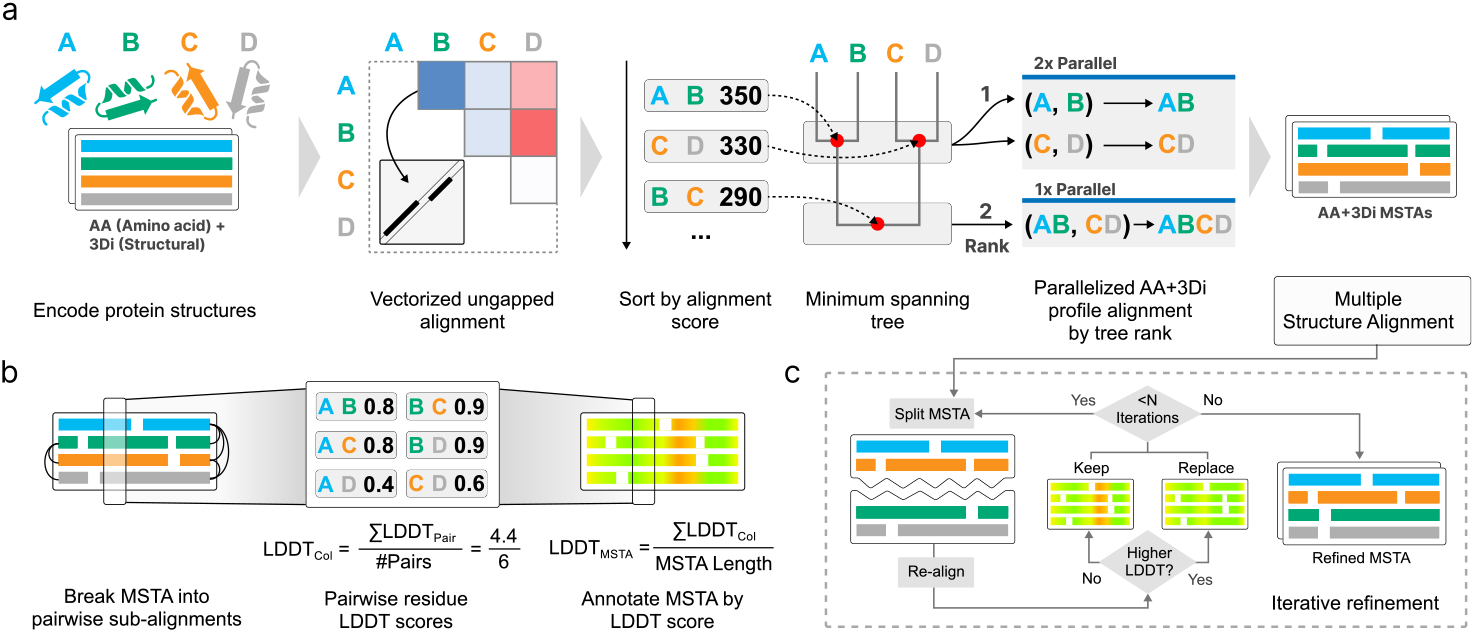
Multiple structure alignment and scoring procedures in FoldMason. (a) MSTA computation with FoldMason. FoldMason represents input protein structures as strings using the 3Di+AA alphabet and computes an ungapped alignment between each pair. Pairs are sorted by alignment score and used to construct a minimum spanning guide tree. Progressive alignment of AA+3Di structure profiles is performed following the guide tree leaf-to-root, with independent alignments computed in parallel based on rank within the guide tree. (b) Computation of confidence scores using LDDT. The input MSTA is broken into pairwise subalignments on which LDDT scores are computed; per-residue LDDT scores are mapped back to their corresponding positions in the MSTA and averaged to give a column LDDT score. Column LDDT scores are averaged to give the final MSTA LDDT score. (c) MSTA refinement by iterative LDDT-maximising. The input MSTA is randomly split into two sub-MSTAs. Resulting gap-only columns are discarded and the sub-MSTAs are re-aligned. At each iteration, the average MSTA LDDT score is computed; the input MSTA is replaced upon improvement to the LDDT score. This is included as an optional procedure after completion of the basic workflow.

Several metrics (e.g., 27, 28) have been developed to assess pairwise superposition-free local similarity. Of these, the local distance difference test (LDDT) is predicted and used by AlphaFold2, thereby becoming a community standard. FoldMason extends LDDT towards MSTAs for a reference-free measure of alignment quality based on conservation of local structural neighbourhoods (**Fig. 1b**). Optionally, FoldMason performs iterative refinement of MSTAs to maximize LDDT (**Fig. 1c**). For easy interpretation of MSTAs, FoldMason can produce interactive visualisations of its results.

We benchmarked FoldMason in comparison to gold-standard structure and sequence aligners on a reference-based Homstrad protein structure dataset and a reference-free set of 1,000 AFDB clusters. We show FoldMason’s scalability on a large AFDB cluster. Finally, we provide case studies of how FoldMason can align flexible protein structures and its use in structure-based phylogeny on Flaviviridae glycoproteins.

We compared FoldMason’s alignments of the Homstrad protein families against those generated by a suite of structure- and sequence-based tools (**Fig. 2a**), including US-align (12), mTM-align (13), MUSTANG (17), Matt (18), Caretta-shape (15), MAFFT (3), MUSCLE5 (4), FAMSA2 (6) and Clustal Omega (5). The Homstrad dataset consists of 1,032 protein structure families with manually curated reference multiple alignments (29). Alignments from structure-based tools exhibited greater accuracy than those from sequence-based tools based on the sum-of-pairs metric (**Fig. 2a**). Structure-based alignments recovered, on average, 6.5% more correctly aligned residue pairs from reference alignments (sensitivity) and possessed a 8.3% greater proportion of aligned residue pairs matching those in the reference alignments (specificity). Notably, US-align, mTM-align, Caretta-shape and FoldMason also include amino acid comparison, thus their alignments represent a blend of sequence and structure data. Caretta-shape (15), performed better than sequence-based methods, but worse than other structure-based methods on this dataset. Structure-based tools were 7.1% more sensitive and 9.5% more specific when excluding Caretta-shape alignments. Importantly, FoldMason reached the accuracy of structure-based aligners (F1 score 0.894, below MUSTANG, US-align and mTM-align; **Table S1)**.

**Fig. 2.**
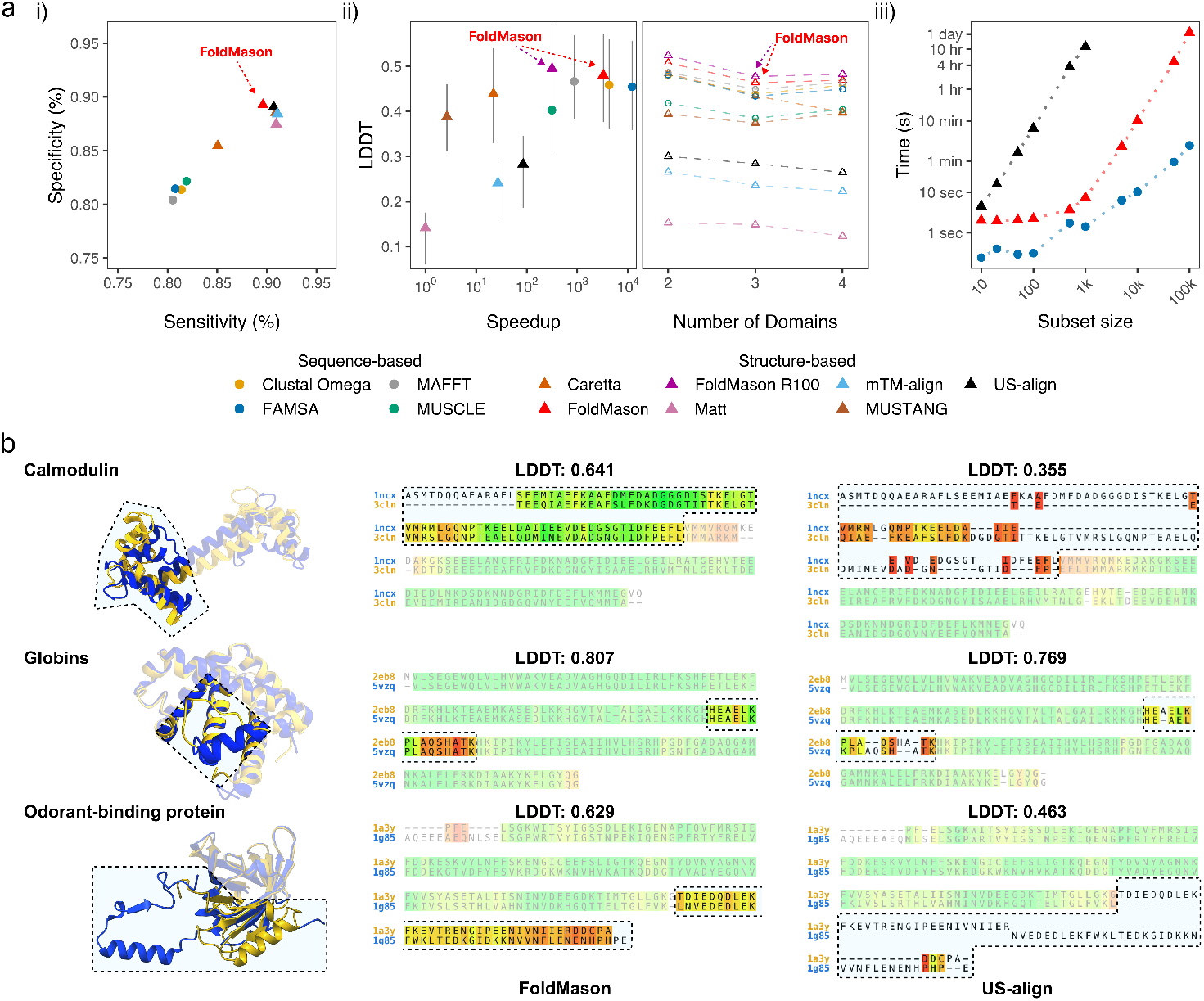
FoldMason’s accuracy and speed compared to other tools on benchmarks. (a) Performance comparison of FoldMason to structure- and sequence-based aligners.i)Sum-of-pairs scores of test alignments compared to Homstrad reference alignments. Sensitivity and specificity are defined as the proportion of aligned residues in the reference found in the test, and from the test in the reference, respectively. ii) Local Distance Difference Test (LDDT) scores of alignments on the 150 AFDB cluster benchmark dataset. Scores are shown with respect to speedup from the slowest tool (left panel) and number of TED domains assigned to each protein. Error bars indicate lower- and upper-quartiles of LDDT scores over all clusters. Speedup per-tool was calculated using the average runtime over the dataset relative to Matt. FoldMason MSTAs with refinement (FoldMason R100) were refined and scored by maximising LDDT over each MSTA for 100 iterations. iii) Run times of sequence- and structure-based tools on varying size subsets of an AFDB cluster (representative UniProt accession A0A258JAY9). Tools are broadly categorised as structure-based or sequence-based. (b) FoldMason and US-align alignments of flexible protein structures. Residues are shaded by the LDDT score of their alignment column.

The manual curation of Homstrad ensures high quality reference alignments but includes mainly small groups of mostly below-average-length, single domain proteins, which do not fully represent most protein families (**Supp. Fig. 1)**. We therefore created a reference-free benchmark dataset based on full-length, high-quality AlphaFold2 predicted structures. For this benchmark, we assessed alignment quality using the average LDDT score, which correlates well with reference-based metrics on the Homstrad dataset (**Supp. Fig. 2)** and can be computed by FoldMason. We leveraged a combined database of AFDB clusters and CATH domain assignments from The Encyclopedia of Domains (TED; 30, 31) to create a benchmark dataset consisting of 150 multidomain protein structure clusters (Online Methods). Each cluster consists of 20 protein structures, containing 2-4 domains, that match at the CATH topology (T) level but are unique at the homologous superfamily (H) level, resulting in a set of highly divergent but globally alignable structures. We then ran the suite of sequence- and structure-based tools on each cluster and computed the average LDDT score for the resulting alignments (**Fig. 2a ii, Table S2)**.

FoldMason exceeded the accuracy of all structure-based tools over this dataset, achieving on average 19.7%, 23.9%, 9.19%, 33.8% and 4.15% higher MSTA LDDT scores than US-align, mTM-align, MUSTANG, Matt and Caretta-shape, respectively. Compared to sequence-based tools, FoldMason achieved 2.16%, 2.55%, 1.36% and 7.77% higher MSTA LDDT scores than Clustal Omega, FAMSA2, MAFFT and MUSCLE, respectively. Notably, FoldMason achieves this while operating at the speed of sequence-based tools, which were on average 307x faster than structure-based tools. FoldMason on average aligned each cluster in under a second, achieving a 3261x speedup when compared to the slowest tool, Matt, while being 4x slower than the fastest tool, FAMSA2. When run with 100 random refinement iterations, FoldMason improved its average MSTA LDDT by 1.47%, with runtime 317x faster than Matt and 38x slower than FAMSA2.

To demonstrate the ability of FoldMason to scale to datasets of this size, we aligned subsets of varying sizes from 10 to 100,000 structures from a large AFDB cluster (representative UniProt accession A0A258JAY9, average length 301.396 amino acids, total 533,657 structures) using US-align (12) and FAMSA2 (6) as representative structure and sequence aligners, respectively (**Fig. 2a iii, Table S3)**. FAMSA2 showed superior scaling performance, able to align 100,000 sequences in 147.49 seconds. On the same set, FoldMason took 92899.10 seconds (∼630x slower), while US-align failed to complete the set of 5000 structures within 2 days. Notably, FoldMason aligned 50,000 structures in under half the time taken for US-align to align 1000 structures. Proteins are flexible and can undergo conformational changes, potentially impacting MSTA methods. Therefore, we tested the ability of structural aligners to handle proteins that exhibit such flexibility (**Fig. 2b**). Calmodulin (CaM) belongs to the EF-hand family of Ca^2+^-sensing proteins (33) and adopts different conformations when bound to calcium and calmodulin-binding enzymes. Calmodulin’s structure consists of N- and C-terminal domains (N-lobe and C-lobe, respectively) connected by a flexible linker domain; once activated by binding calcium, it adopts many conformations ranging from fully extended to collapsed with the linker wrapping around a bound target protein (34). This flexibility is present in many CaM structures deposited on the Protein Data Bank (PDB) and poses problems for current structure alignment tools.

We identified two CaM structures (PDB identifiers 1ncx, 3cln) in which flexibility causes the N- and C-lobes to be non-superposable; FoldMason successfully aligns the N-lobes of both structures, whereas US-align, a superposition-based method, fails to do so (**Fig. 2b**, top row). Another example of internal flexibility hindering structural alignment came from structures of apo- and nitric oxide-bound myoglobins (PDB identifiers 2eb8 and 5vzq, respectively) from sperm whales (35). Myoglobins are heme-containing proteins that facilitate the diffusion of oxygen in the muscle (36). Despite near-identical amino acid sequence, flexibility of a loop inside the structure of the apo-bound myoglobin proved problematic for the superposition-based alignment of US-align; FoldMason however successfully aligned all residues. Finally, we compared odorant-binding proteins (OBP) found in pigs and cows (PDB identifiers 1a3y and 1g85, respectively). Whereas the pig OBP is a monomer with a helix and strand of sheet in its C-terminus, the cow OBP is a dimer in which the C-terminal domain of each constituent monomer have been swapped (17). While the pig OBP monomer mostly matches one of the cow OBP monomers, the swapped domain is oriented differently and is an issue for the superposition-based US-align, but is successfully recovered by the FoldMason.

While both FoldMason and US-align compute MSTAs, the comparisons in this section were performed pairwise to better illustrate the differences between the tools on structurally flexible regions. We additionally built MSTAs of each protein using each structural aligner and compared their average LDDT scores (**Table S4)**. FoldMason alignments consistently achieved the highest average LDDT across each protein family, with the exception of globins, where Caretta-shape matched its performance in pairwise alignment, and achieved 2.7% higher in MSTA. FoldMason achieved 22.3%, 16.6% and 3.8% higher average MSTA LDDT scores than US-align for calmodulin, OBP and globins, respectively. These cases suggest that FoldMason is more tolerant of structural flexibility compared to superposition-based structural aligners, as it depends on local interactions only.

Phylogenetic analysis commonly relies on MSAs, here MSTAs can extend these analyses to proteins related beyond the twilight zone. There have been multiple studies applying Foldseek and its 3Di alphabet for remote phylogenetic analysis. For instance, Puente-Lelievre et al. (32) demonstrated that MSTAs of 3Di sequences offer information complementary to traditional sequence data, thereby aiding phylogenetic reconstruction. Additionally, Moi et al. (37) introduced FoldTree for the comparison of distantly related proteins, using Fold-seek to compute trees from a sequence identity matrix generated by structural pairwise alignments.

Mifsud et al. (38) investigated the distant taxonomic relationships between viral glycoproteins, proteins that were not alignable through sequence-based methods, by assembling a MSTAs from Foldseek pairwise alignments. FoldMason simplifies by directly generating a MSTAs and therefore eliminating the need for ad-hoc approaches. To demonstrate FoldMason’s capability in this context, we reconstructed the phylogenies of the E, E1, and E2 glycoproteins reported by Mifsud et al. (see Figure 3). The topology of the trees generated from FoldMason MSTAs generally matches those reported, with all annotated clades successfully recovered.

**Fig. 3.**
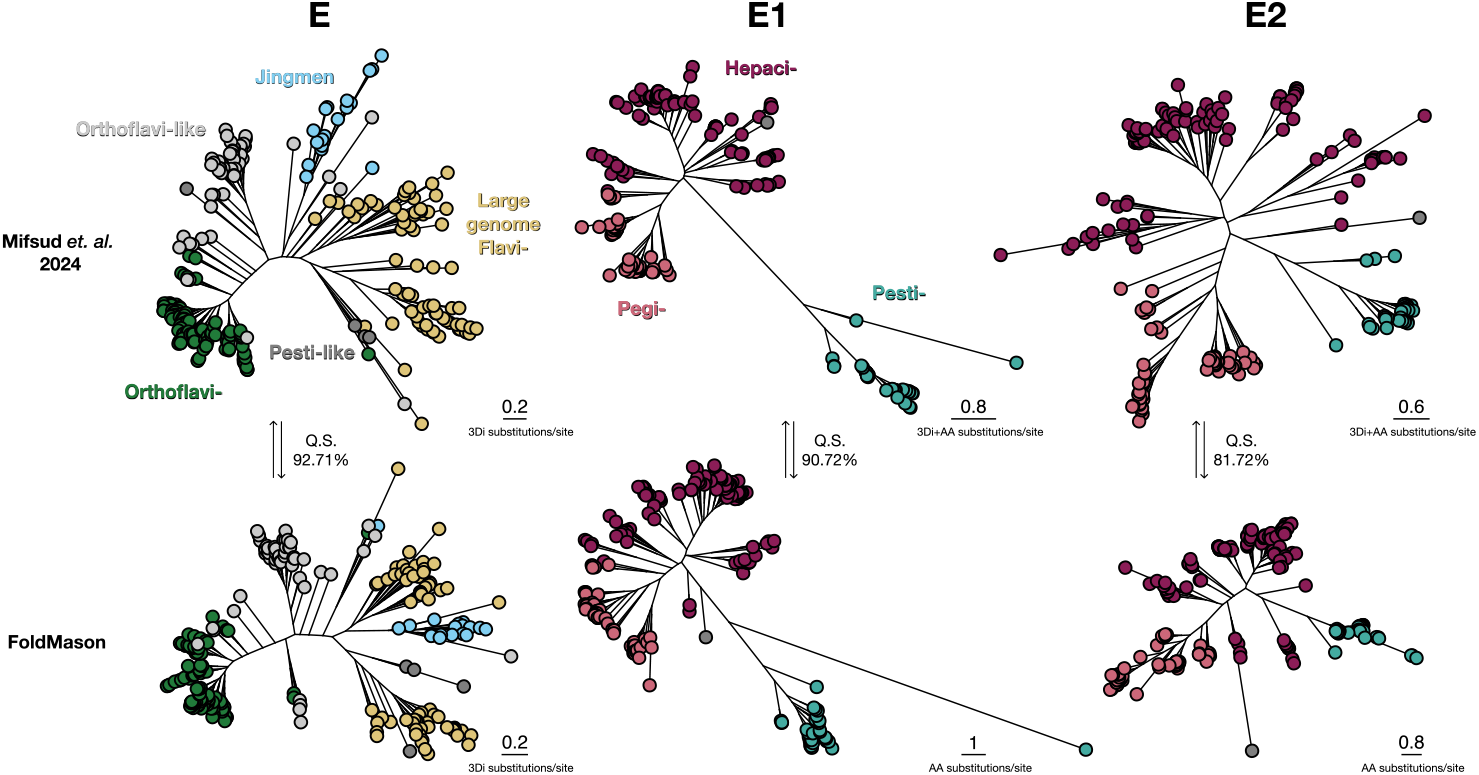
FoldMason’s directly computed MSTA supports structural phylogenies previously inferred by manual MSTA reconstruction. Mifsud et al. followed the procedure described in (32) to build MSTAs of the viral E, E1 and E2 glycoproteins and infer their structural phylogenies (top). We aligned the reference structures from Mifsud et al. using FoldMason (Online Methods) and used the MSTAs for phylogeny inference (bottom). Congruence between the Mifsud et al. and FoldMason-derived phylogenies is high, as indicated by their quartet similarities (Q.S). Q.S. indicates quartet similarity between the corresponding Mifsud et. al. and FoldMason-derived phylogenies.

This congruence is reflected by the high quartet similarity between the reported trees and those constructed from FoldMason; 92.71%, 90.72%, and 81.72% for E, E1, and E2, respectively. We observed comparable bootstrap support across all nodes in the FoldMason trees (87.3-90.2%) and those from Mifsud et al. (87.1-93.3%). When only considering corresponding nodes (85/245 nodes, 109/188 nodes and 100/187 nodes for E, E1 and E2 respectively), the difference was negligible (difference range 1.2-1.6%), indicating FoldMason MSTAs can produce trees of equal quality. Overall, FoldMason through its high quality alignments as well as the access to the 3Di information, makes it a powerful tool for phylogenetic analysis beyond the twilight zone.

For future work, we plan to integrate the ProstT5 protein language model (39) to directly predict 3Di from amino acid sequences, eliminating the need for slow structure predictions. This will accelerate input generation for FoldMason by over 3000× compared to optimized ColabFold prediction and would be particularly beneficial for studies involving long proteins, as in Mifsud et al. Moreover, we plan to tightly integrate FoldMason within the Foldseek webserver, allowing users to generate MSTAs directly from Foldseek search results, and extend FoldMason to protein complexes through the integration of Foldseek-Multimer (40). Thus, turning FoldMason into a comprehensive suite to study structures and evolution across the tree of life.

In summary, FoldMason reaches or exceeds alignment accuracy of state-of-the-art structure-based aligners while operating several orders of magnitudes faster, scaling to large datasets, and offering user-friendly visualisations and web-server (**Fig. 4)** making it an indispensable tool in the post-AlphaFold era.

**Fig. 4.**
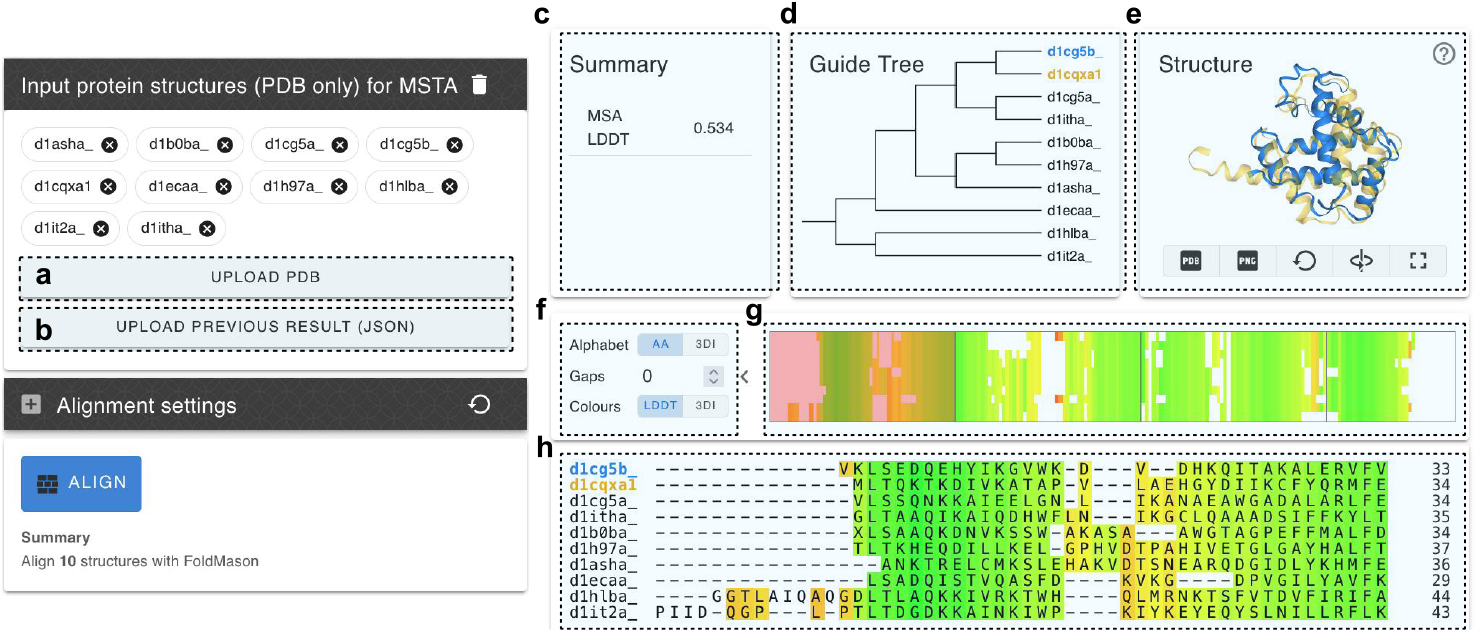
Interactive HTML MSTA visualisation in the FoldMason webserver. Launch page (left side): users can (a) upload protein structures in PDB format (mmCIF support soon to follow) as well as (b) previous FoldMason results in JSON format. Results page (right side): (c) MSTA LDDT score and metadata related to the FoldMason run. (d) Guide tree used in the FoldMason alignment; only shown when generated with the basic workflow. (e) Interactive structure viewer. MSTA structures can be selected for examination against a selected ‘representative’ structure; the representative is initially the first structure in the MSTA but can be changed by the user. Each selected sequence (shown in yellow) is superposed on to the representative (shown in blue) with TM-align using their induced PSA from the MSTA. Users can generate PDB files with superposed structures and screenshots of the current state within the viewer. (f) Adjustable parameters for the MSTA view. Users can toggle between AA/3Di alphabets, set a percentage gap threshold to hide gappy columns and change between LDDT-based and 3Di-based colorschemes. (g) A ‘minimap’ allowing users to jump between aligned blocks within the MSTA view. The currently visible block is highlighted in red; clicking another segment in the minimap will scroll the view to the corresponding block. (h) The MSTA view. By default, column background colors correspond to per-column LDDT values calculated by FoldMason. Users can adjust the structures shown in the structure viewer panel (e) directly from the MSTA view. Left-clicking a structure name aligns that structure to the current representative and add it to the viewer; clicking again removes the structure. A new representative structure can be selected by alt-clicking a structure name; all other currently-selected structures are automatically aligned to the new representative.

## SUPPLEMENTARY MATERIALS

### Algorithm: Using Foldseek as a library

FoldMason is a standalone command-line software that builds upon Foldseek’s functionality by integrating it as an internal library. In the following, we describe FoldMason’s algorithm, indicating where Foldseek’s modules are used.

### Algorithm: Database creation

Protein 3D structures are discretized to 1D using the 3D interactions (3Di) alphabet described previously (22). FoldMason makes use of amino acid (AA), 3Di and *α* Carbon backbone coordinates extracted from input protein structures and stores these as Foldseek databases.

### Algorithm: Guide tree

To generate a guide tree, FoldMason first computes an all-vs.-all comparison of the input protein structures using a SIMD-accelerated gapless alignment algorithm implemented in Fold-seek. A minimum-spanning tree approach is taken in order to generate the guide tree for the alignment. Briefly, all pairwise alignments are sorted by their score in best-to-worst order. We then iteratively select the best scoring pairwise alignment as the next node in the guide tree. All subsequent pairwise alignments containing a structure from the new node are then updated to reflect the newly created node. This selection process is repeated until the full guide tree has been constructed. Finally, we re-order the tree to maximise the number of alignments that can be performed in parallel. This reordering marks sets, termed *indAlnSets*, where all alignments within a set can be computed independently.

### Algorithm: Progressive Alignment

We utilise a progressive algorithm to align structures along the generated guide tree. Alignment is performed in rounds as determined by the parallelisation-maximizing reordering. In the first round, all *indAlnSets* (in this case, a single set, containing all leaf nodes) are globally aligned using both TM-align and a semi-global implementation of the Gotoh algorithm (41) with affine gap costs, aligning 3Di and amino acid sequences jointly. The alignment with the highest LDDT (see below) is kept and a bitpacked representation of each sequence is updated to reflect the current alignment status.

We implemented a bitpacked data structure in which 8 bits are used to represent each sequence state, to reduce memory usage, particularly in gappy alignments. The first bit indicates whether the remaining 7 bits represent sequence (0) or gap (1) states; the 7 bits then store either an ASCII character corresponding to a AA/3Di residue or a gap count (maximum 127), respectively. This allows for a series of 127 gaps to be represented as a single byte. Position-specific score matrices (PSSMs) for both AA and 3Di sequences are created to represent the aligned sequences. In subsequent rounds, either sequence-PSSM or consensus-PSSM alignments are performed using the Gotoh algorithm. In consensus-PSSM alignment, extraction of the consensus sequence is done on the PSSM with the lower entropy (*N*_eff_).

Bitpacked representations are again updated to reflect the new alignments, and new PSSMs are created (e.g. a single PSSM may represent ≥ 4 structures after PSSM-PSSM alignment). This is repeated until the guide tree has been fully traversed. All alignments in each round are performed in parallel or sequentially if there are insufficient CPU cores.

### Algorithm: Computation of average MSA LDDT scores using **msa2lddt**

The Local Distance Difference Test (LDDT) is a superposition-free metric for assessing the similarity of protein structures based on their local structural neighbourhoods (27). Briefly, calculation of the LDDT score of equivalent residues in two structures involves 1) collecting distances between pairs of atoms falling within some inclusion radius of each residue (typically 15Å); 2) calculating the fraction of distances which are preserved in the atomic neighbourhoods of both residues over a set of distance thresholds (0.5Å, 1Å, 2Å and 4Å); and 3) taking the average of these fractions to calculate the final LDDT score.

We implemented a module in FoldMason, `msa2lddt`, in which we adapted the LDDT for use as a reference-free MSA quality metric. As input, it accepts a FASTA-format MSA file and associated structures. `msa2lddt` is not specific to Fold-seek’s output, it accept any MSAs as long as individual entry accessions can be mapped to structures. It produces either a user-friendly interactive HTML report of the given MSA (see below), or machine-readable data in JSON format for downstream use.

The average MSA LDDT score is computed as follows: For all pairs on structures in the MSTA, we compute the LDDT score of aligned residues in match columns; columns with gaps are discarded. A per-column LDDT score is computed by taking the sum of LDDT scores over all PSAs for a given column, averaged over the number of residue pairs contributing to that sum. The MSA LDDT score is the average of the per-column LDDT scores over the length of the alignment.

Users can optionally specify the maximum proportion of gaps within a column for it to contribute to the MSA LDDT score (pair threshold); by default, all per-column LDDT scores are included. Specifying 100%, for instance, ensures only gapless columns will be included, which has the effect of only computing the LDDT score of the ‘conserved core’ of all structures.

### Algorithm: Iterative MSA refinement using refinemsa

We implemented an iterative alignment refinement procedure based on randomized partitioning (42) maximising LDDT score in the refinemsa module of FoldMason. This module takes as input a FASTA-formatted MSA and associated structure. As before, the input MSTA can be generated by any tool, if structures can be associated.

Refinement can be optionally enabled during regular MSA generation with the `easy-msa` workflow, or as its own separate module. Due to computational cost, it is disabled by default.

### Algorithm: Software output

The `easy-msa` module is the main entry point for FoldMason users. It accepts PDB/mmCIF structure files and generates two FASTA-formatted MSA files, both sharing the same underlying alignment, one containing AA sequence and the other 3Di. Optionally, it can save the guide tree used to guide the progressive alignment in Newick-format, calculate the LDDT score of a given MSA and generate an interactive HTML visualisation (described above).

### Algorithm: Interactive HTML visualisation

An interactive visualisation can be requested through the command-line interface or using the webserver. The web-server is a further development of the MMseqs2 server (43). The MSTA result visualization is structured in three main tiers; briefly, from top to bottom 1) information panel, 2) a control panel, and 3) the MSA view panel. The information panel consists of either two or three panels when generated by `msa2lddt` or `easy-msa`, respectively. The left-most panel provides runtime information (command string, input file paths, etc) and statistics of the alignment. If generated by `easy-msa`, a middle pane is inserted which displays the guide tree used to guide the progressive alignment in FoldMason. The right-most panel contains a 3D protein structure visualisation. By default, it shows the two top-most proteins from the MSA superposed using TM-align (14) via the induced pairwise alignment from the MSA; one protein (in blue) is chosen as the reference on to which other target structures (in yellow) are superposed. Users can select a different entry as reference and additional proteins to be superposed. The control panel consists of toggleable visualisation settings and a ‘minimap’ for MSA navigation. In the toggle settings panel, from top to bottom, the user is able to adjust: 1) the sequence alphabet, 2) the column-gappiness percentage threshold, 3) the MSA colorscheme. The alphabet can either be set to amino acid (default) or the 3Di alphabet. The gappiness percentage controls which columns are shown or hidden based on the proportion of gaps in each column; for instance, if set to 30%, columns containing less than ≤30% sequence characters will be hidden from the view. Two colorschemes can be selected: by default, a divergent red-to-green scheme based on the per-column LDDT scores is shown (red 0%, green 100%). Alternatively, a 3Di state-based colorscheme, can be selected (32). The ‘minimap’ provides an easy way to navigate the MSA by displaying clickable MSA blocks corresponding to those in the MSA view; if the user clicks one block, the view will automatically scroll them to the corresponding position in the MSA. Finally, the MSA view panel displays the MSA in CLUSTAL format with sequence accessions on the left, sequence in the middle, and sequence positions at end of rows (not including gaps) shown on the right. The user can select additional structures from the MSA to visualise and superpose in the structure viewer by left-clicking the corresponding sequence header in the MSA view. Alt-clicking on the reference structure will clear all structure selections, and on any other structure will result in this structure being chosen as the new reference structure.

### Benchmarks

We benchmarked FoldMason against a variety of structure and sequence aligners: mTM-align (13), US-align (), MUS-TANG (17), Matt (18) and Caretta-shape (15) were chosen as representative structure aligners, whereas Clustal Omega (5), MUSCLE 5 (4), MAFFT (3) and FAMSA 2 (6) were chosen as representative sequence aligners. We tested FoldMason commit 92b1690, Clustal Omega v1.2.4, FAMSA v2.2.2-7eb7612, MAFFT v7.520, Matt v1.0, mTM-align v20220104, US-align v20240730, MUSCLE v5.1, MUSTANG v3.2.4 and Caretta-shape commit 998125b. All tools were executed with default parameters on a server with 1x AMD EPYC 7702P 64-core CPU and 1,024 GB RAM memory. For the Homstrad and AFDB cluster benchmarks, all tools were run single-threaded. In the AFDB scaling benchmark, FoldMason and FAMSA were run with 64 cores, and US-align, which does not support multi-threading, was run single-threaded.

### Benchmarks: Homstrad

The Homstrad database (44) consists of 1032 protein families with manually curated reference alignments. Reference alignments and the corresponding PDB files were obtained from the May 2024 release (Data availability). A cleaning script `clean_homstrad.py` (Code availability) was used to generate PDB-format protein structure files on which structure-based tools were run, as well as to generate AA sequence files for the sequence-based aligners. We ran each tool on the entire set of Homstrad protein families. We computed the sum-of-pairs (SP), total columns score (TC) and column score (CS) using the aln_compare module in T-Coffee (45). We computed the SP score in two directions: 1) using the Homstrad reference MSA as the reference alignment in aln_compare (-al1), and 2) using a test MSA generated by one of the tools as the reference alignment. In the former direction, we identify pairs of aligned residues from the reference MSA in each test MSA, approximating a sensitivity measure; and in the latter, residue pairs from the test MSA are found in the reference MSA, approximating a precision measure. In addition, we computed the LDDT score of all MSAs, including the Homstrad reference MSAs, using the `msa2lddt` module. We used the structure database used during the FoldMason alignment during this calculation.

### Benchmarks: AFDB cluster database

Protein structure clusters were selected from the clustered AlphaFold database we described previously (23). AFDB Cluster data first release were retrieved (Data availability) and a SQLite3 database was built. In addition, we integrated domain assignments for each structure from The Encyclopedia of Domains (TED; 46). For each TED domain annotation, we took the corresponding CATH (47) classification and stored it in the database; multidomain proteins with ≥ 2 TED annotations are thus annotated with ≥ 2 CATH classifications. In the case of single TED annotations with multiple corresponding CATH classifications, we selected only the first CATH classification. From this combined database, we selected clusters in which the domain architecture of member structures, i.e. the linear arrangement of their assigned CATH domains, matched that of the corresponding cluster representative structure up to the topology (T) level but were unique at the homologous superfamily (H) level. We took a total of 150 clusters to use as a benchmark dataset, consisting of 50 two-domain clusters, 50 three-domain clusters and 50 four-domain protein clusters. These clusters were further filtered to contain only the first 20 members inclusive of the representative structure. We ran each tool on the AFDB clusters, measuring LDDT score of each MSA using `msa2lddt`. We measured the run time of each tool over the entire set of clusters; speedup for each tool was computed in comparison to Matt, which was the slowest tool tested. FoldMason alignments with refinement were computed using the refinemsa module with 100 iterations with fixed seed 48335597; refinement was performed maximising LDDT scores over the full MSTAs inclusive of gappy columns (shown as FoldMason R100 in **Fig. 2a ii**) left and right panels, respectively).

### Benchmarks: Flexibility

We selected three example protein families exhibiting structural flexibility to test FoldMason’s ability to construct MSAs in various scenarios. US-align performed the best in out of structural aligners in the previous benchmarks and was thus used here to compare; however, we also constructed alignments using other structural aligners. Example protein structures for calmodulin (PDB identifiers 1jfj, 1ncx, 2sas and 3cln), globins (PDB identifiers 1h97, 1hho, 1mbn, 1oj6, 1urv, 2eb8, 2hhb, 2oif and 5vzq) and odorant-binding proteins (PDB identifiers 1a3y, 1g85, 3fiq, 3zq3 and 5ngh) were retrieved from the RCSB Protein Data Bank (48, Data availability). We constructed MSTAs using FoldMason, US-align (), MUSTANG (17), Matt (18) and Caretta-shape (15). Matt optionally produces ‘bent’ alignments, where structurally impossible bends and breaks are permitted; we included these in our analysis. LDDT scores of both pairwise alignments and MSTAs were computed using the `msa2lddt` module in Fold-Mason. Structures in pairwise alignments were superposed and visualised using UCSF ChimeraX 1.6.1 (49).

### Benchmarks: Scalability

In order to test the ability of FoldMason to align large structure datasets, we queried our AlphaFold2 cluster database for clusters with average length >300 amino acids. The cluster with representative structure A0A258JAY9 was the largest cluster identified, containing 533,657 members with average length 301.396 and average pLDDT 89.099, and was thus chosen for this benchmark. The cluster was then subset at several sizes (10, 20, 50, 100, 200, 500, 1000, 5000, 10000, 50000, 100000) to create benchmark datasets of differing size. We chose US-align and FAMSA 2 as representative structure and sequence aligners, respectively. We ran each tool on each size subset; FAMSA2 and FoldMason successfully completed all sets, whereas US-align was stopped after 214 hours on the 500 structure subset.

### Benchmarks: Glycoprotein Phylogeny

Data from (38) was retrieved from Zenodo (50, Data availability, and private communications), including protein structures, alignments and phylogenetic trees. Reference trees for E, E1 and E2 glycoproteins were selected as described in (51): 3Di for E, and partitioned 3Di and AA trees for E1 and E2, trimmed to remove any columns containing ≥35 gaps. E, E1 and E2 protein structures were aligned using FoldMason with default settings and then trimmed using trimAl (52, v1.4.rev15 build[2013-12-17]) with a gap threshold of 35% as in (38). For the E family, the 3Di character alignment was used with the 3Di substitution matrix used in (32). For E1/2, the best ranked AA substitution model was determined using ModelFinder (53); EX_EHO+R6 and WAG+F+R6 substitution models were used for E1 and E2, respectively. Phylogenetic trees were built using IQ-Tree v2.3.0 (54) with 1000 ultrafast bootstrap replicates (55). Trees were visualised using ggtree v3.12 (56). Quartet similarity between FoldMason trees and reference trees from (38) was computed using the tqDist algorithm in the Quartet R library v1.2.6 (57, 58).

## Data and materials availability

Homstrad: https://homstrad.mizuguchilab.org/homstrad/data/ AFDB clusters: https://afdb-cluster.steineggerlab.workers.dev/ Glycoprotein data: https://zenodo.org/doi/10.5281/zenodo.10616318 PDB: files.wwpdb.org/pub/pdb/data/structures/all FoldMason source code and pre-built binaries are freely available under the GPL-3.0 license. Source code and binaries for FoldMason can be found at github.com/steineggerlab/foldmason. Analysis scripts used in this paper are available at github.com/steineggerlab/foldmason-analysis. The webserver is available at search.foldseek.com/foldmason, with source code available at github.com/soedinglab/mmseqs2-app.

## Acknowledgments

We thank Cedric Notredame, Adam Gudyś and Sebastian De-orowicz for their insightful discussions, and Eli Levy Karin from ELKMO for the valuable scientific feedback and careful review and revision of the paper. We thank the EMBL-EBI for releasing the “EMBL-EBI Icon fonts for the life sciences” that are used in the webserver. M.S. acknowledges support by the National Research Foundation of Korea grants (2020M3-A9G7-103933, 2021-R1C1-C102065, 2021-M3A9-I4021220 and RS-2024-00396026), Samsung DS research fund, Creative-Pioneering Researchers Program and AI-Bio Research Grant through Seoul National University. M.M. acknowledges support from the National Research Foundation of Korea (grant RS-2023-00250470).

## Author contributions

C.L.M.G., M.M. and M.S. designed the FoldMason algorithm, software and webserver. C.L.M.G. designed the figures, performed the benchmarks and wrote the manuscript, with contributions from all authors.

**Supplementary Figure 1.**
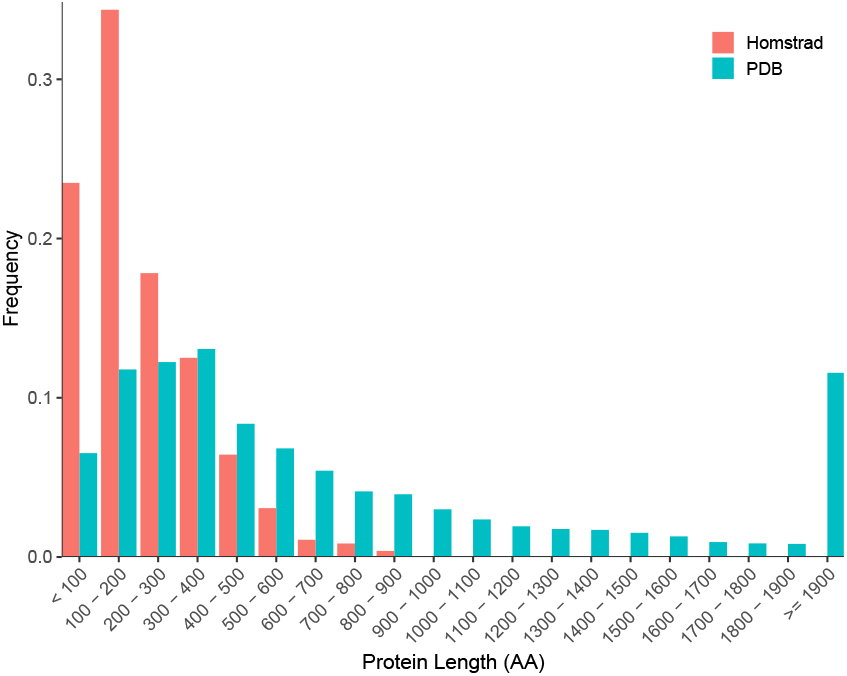
Protein length (AA residue) distributions of Protein Data Bank (PDB) and Homstrad databases. Binned PDB protein sizes (N=222624) were obtained from the PDB website (www.rcsb.org/stats/distribution-residue-count; accessed 2024-07-17). Homstrad protein sizes (N=3422) were binned according to the ranges in the PDB data. Length ranges are displayed as their frequency in their respective dataset.

**Supplementary Figure 2.**
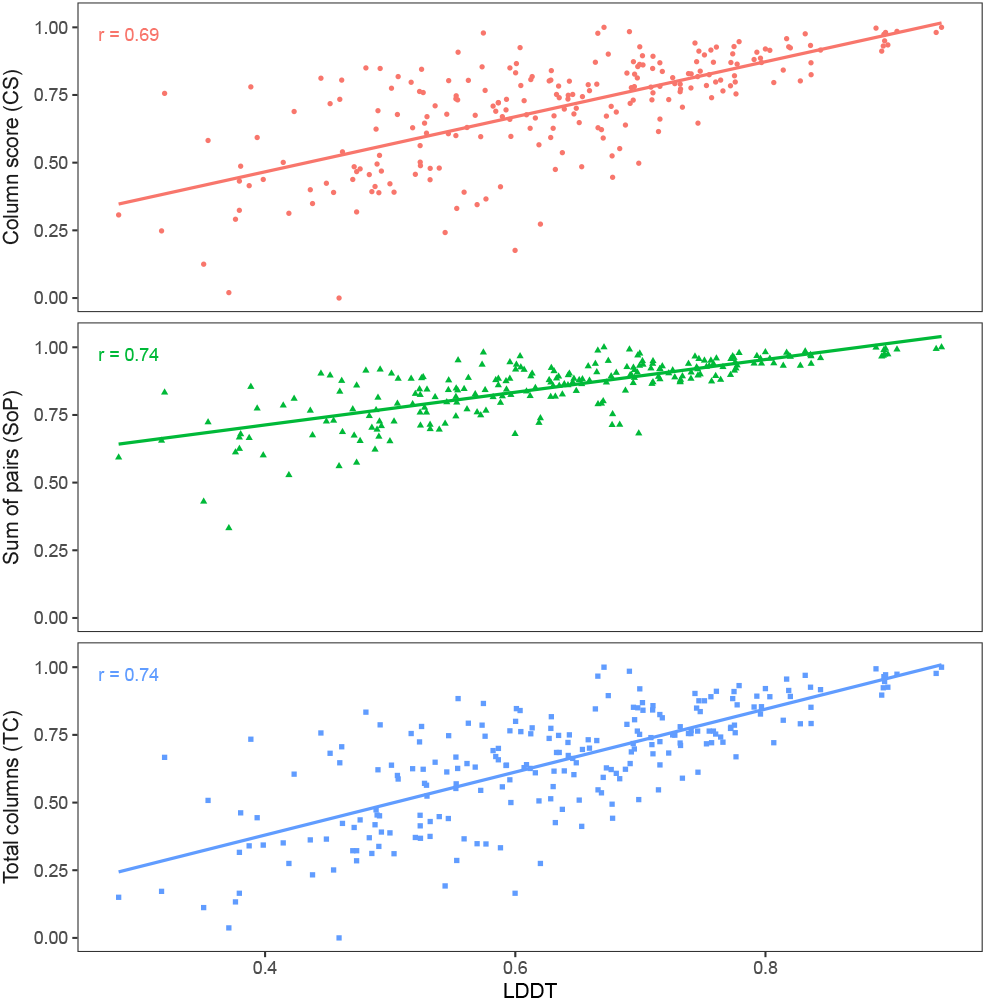
Correlation of alignment LDDT score and reference-based alignment quality metrics on Homstrad families with ≥**4 members**. Reference-based scores (sum of pairs, total columns score, column score) were computed for FoldMason alignments against reference alignments from the Homstrad database using the aln_compare module in T-Coffee. LDDT scores were computed using the `msa2lddt` module from FoldMason. All measures correlate highly (Pearson’s *r*).

**Supplementary Table 1.** Performance of aligners on the Homstrad benchmark dataset. Values given are sum-of-pairs scores computed between test alignments and Homstrad reference alignments. Sensitivity and specificity are defined as the proportion of aligned residues in the reference found in the test (i.e. calculating sum-of-pairs using Homstrad alignment as reference), and from the test in the reference, respectively. F1 score was computed from the sensitivity and specificity scores shown.

| Method | Sensitivity | Specificity | F1 Score |
| --- | --- | --- | --- |
| MAFFT | 0.804 | 0.805 | 0.805 |
| FAMSA | 0.814 | 0.808 | 0.811 |
| Clustal Omega | 0.814 | 0.814 | 0.814 |
| MUSCLE | 0.821 | 0.819 | 0.820 |
| Caretta-shape | 0.855 | 0.851 | 0.853 |
| Matt | 0.875 | 0.910 | 0.892 |
| FoldMason | **0.892** | 0.896 | 0.894 |
| MUSTANG | 0.885 | 0.909 | 0.897 |
| mTM-align | 0.884 | **0.911** | 0.897 |
| US-align | 0.891 | 0.907 | **0.899** |

**Supplementary Table 2.** Performance of aligners on the AFDB clusters benchmark dataset. Average LDDT values were computed using FoldMason `msa2lddt` over all, 2-domain only, 3-domain only and 4-domain only MSTAs. FoldMason R100 refers to FoldMason MSTAs built with 100 random refinement iterations. Runtime (s) values are averaged over all alignments in the dataset.

| Tool | Average MSTA LDDT |  |  |  | Average runtime (s) |
| --- | --- | --- | --- | --- | --- |
|  | All | 2-domain | 3-domain | 4-domain |  |
| Caretta-shape | 0.439 | 0.484 | 0.434 | 0.398 | 140.0 |
| Clustal Omega | 0.459 | 0.479 | 0.439 | 0.458 | 0.729 |
| FAMSA | 0.455 | 0.481 | 0.434 | 0.449 | **0.256** |
| FoldMason | 0.480 | 0.508 | 0.464 | 0.469 | 0.947 |
| FoldMason R100 | **0.495** | **0.524** | **0.477** | **0.483** | 9.74 |
| MAFFT-LINSI | 0.467 | 0.487 | 0.449 | 0.464 | 3.55 |
| Matt | 0.142 | 0.153 | 0.149 | 0.123 | 3089.0 |
| mTM-align | 0.241 | 0.266 | 0.236 | 0.222 | 114.0 |
| US-align | 0.283 | 0.300 | 0.284 | 0.265 | 36.4 |
| MUSCLE | 0.403 | 0.418 | 0.385 | 0.404 | 9.81 |
| MUSTANG | 0.388 | 0.394 | 0.375 | 0.396 | 1168.0 |

**Supplementary Table 3.** Average runtime (s) of FoldMason, FAMSA2 and US-align on the AFDB cluster scaling benchmark dataset. MSTAs were constructed from various sized subsets of a large AFDB cluster (representative UniProt accession A0A258JAY9, average length 301.396 amino acids, total 533,657 structures). N.A. indicates that the alignment did not complete.

| Subset size (Structures) | Runtime (s) |  |  |
| --- | --- | --- | --- |
|  | FAMSA2 | FoldMason | US-align |
| 10 | 0.24 | 2.02 | 4.62 |
| 20 | 0.40 | 1.96 | 15.85 |
| 50 | 0.29 | 2.09 | 98.02 |
| 100 | 0.31 | 2.26 | 394.6 |
| 500 | 1.75 | 3.72 | 12887 |
| 1000 | 1.43 | 7.32 | 41426 |
| 5000 | 6.35 | 138.51 | N.A. |
| 10000 | 10.23 | 608.06 | N.A. |
| 50000 | 57.13 | 17610.18 | N.A. |
| 100000 | 147.49 | 92899.10 | N.A. |

**Supplementary Table 4.**
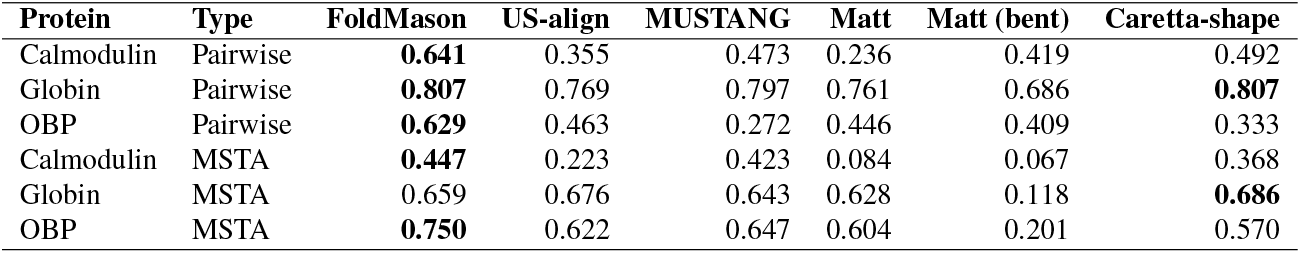
Average LDDT of MSTAs computed by structure aligners on flexible proteins. Calmodulin MSTAs are built from PDB identifiers 1jfj, 1ncx, 2sas and 3cln; pairwise alignments from 1ncx and 3cln. Globin MSTAs are built from PDB identifiers 1h97, 1hho, 1mbn, 1oj6, 1urv, 2eb8, 2hhb, 2oif and 5vzq; pairwise alignments from 2eb8 and 5vzq. Odorant-binding protein (OBP) MSTAs are built from PDB identifiers 1a3y, 1g85, 3fiq, 3zq3 and 5ngh; pairwise alignments from 1a3y and 1g85.

| Protein | Type | FoldMason | US-align | MUSTANG | Matt | Matt (bent) | Caretta-shape |
| --- | --- | --- | --- | --- | --- | --- | --- |
| Calmodulin | Pairwise | **0.641** | 0.355 | 0.473 | 0.236 | 0.419 | 0.492 |
| Globin | Pairwise | **0.807** | 0.769 | 0.797 | 0.761 | 0.686 | **0.807** |
| OBP | Pairwise | **0.629** | 0.463 | 0.272 | 0.446 | 0.409 | 0.333 |
| Calmodulin | MSTA | **0.447** | 0.223 | 0.423 | 0.084 | 0.067 | 0.368 |
| Globin | MSTA | 0.659 | 0.676 | 0.643 | 0.628 | 0.118 | **0.686** |
| OBP | MSTA | **0.750** | 0.622 | 0.647 | 0.604 | 0.201 | 0.570 |

## References

1. Wang, L. & Jiang, T. On the Complexity of Multiple Sequence Alignment. Journal of Computational Biology 1, 337–348 (1994).

2. Feng, D.-F. & Doolittle, R. F. Progressive sequence alignment as a prerequisitetto correct phylogenetic trees. Journal of Molecular Evolution 25, 351–360 (1987).

3. Katoh, K. & Standley, D. M. MAFFT Multiple Sequence Alignment Software Version 7: Improvements in Performance and Usability. Molecular Biology and Evolution 30, 772–780 (2013).

4. Edgar, R. C. Muscle5: High-accuracy alignment ensembles enable unbiased assessments of sequence homology and phylogeny. Nature Communications 13, 6968 (2022).

5. Sievers, F. et al. Fast, scalable generation of high-quality protein multiple sequence alignments using clustal omega. Molecular Systems Biology 7, 539 (2011).

6. Deorowicz, S., Debudaj-Grabysz, A. & Gudyś, A. FAMSA: Fast and accurate multiple sequence alignment of huge protein families. Scientific Reports 6, 33964 (2016).

7. Carpentier, M. & Chomilier, J. Protein multiple alignments: sequence-based versus structure-based programs. Bioinformatics 35, 3970–3980 (2019).

8. Lesk, A. M. & Konagurthu, A. S. Protein structure prediction improves the quality of amino-acid sequence alignment. Proteins: Structure, Function, and Bioinformatics 90, 2144–2147 (2022).

9. Rajapaksa, S., Konagurthu, A. S. & Lesk, A. M. Sequence and structure alignments in post-AlphaFold era. Current Opinion in Structural Biology 79, 102539 (2023).

10. Rost, B. Twilight zone of protein sequence alignments. Protein Engineering, Design and Selection 12, 85–94 (1999).

11. Illergård, K., Ardell, D. H. & Elofsson, A. Structure is three to ten times more conserved than sequence—A study of structural response in protein cores. Proteins: Structure, Function, and Bioinformatics 77, 499–508 (2009).

12. Zhang, C., Shine, M., Pyle, A. M. & Zhang, Y. Us-align: universal structure alignments of proteins, nucleic acids, and macromolecular complexes. Nature Methods 19, 1109–1115 (2022).

13. Dong, R., Peng, Z., Zhang, Y. & Yang, J. mTM-align: an algorithm for fast and accurate multiple protein structure alignment. Bioinformatics 34, 1719–1725 (2018).

14. Zhang, Y. & Skolnick, J. TM-align: a protein structure alignment algorithm based on the TM-score. Nucleic Acids Research 33, 2302–2309 (2005).

15. Akdel, M., Durairaj, J., de Ridder, D. & van Dijk, A. D. J. Caretta – A multiple protein structure alignment and feature extraction suite. Computational and Structural Biotechnology Journal 18, 981–992 (2020).

16. Durairaj, J., Akdel, M., Ridder, D.d. & Dijk, A. D. v. Fast and adaptive protein structure representations for machine learning. bioRxiv 2021.04.07.438777 (2021).

17. Konagurthu, A. S., Whisstock, J. C., Stuckey, P. J. & Lesk, A. M. MUSTANG: A multiple structural alignment algorithm. Proteins: Structure, Function, and Bioinformatics 64, 559–574 (2006).

18. Menke, M., Berger, B. & Cowen, L. Matt: Local Flexibility Aids Protein Multiple Structure Alignment. PLOS Computational Biology 4, e10 (2008).

19. Jumper, J. et al. Highly accurate protein structure prediction with AlphaFold. Nature 596, 583–589 (2021).

20. Lin, Z. et al. Evolutionary-scale prediction of atomic-level protein structure with a language model. Science 379, 1123–1130 (2023).

21. Varadi, M. et al. AlphaFold protein structure database in 2024: providing structure coverage for over 214 million protein sequences. Nucleic Acids Res. 52, D368–D375 (2024).

22. van Kempen, M. et al. Fast and accurate protein structure search with Foldseek. Nature Biotechnology 42, 243–246 (2024).

23. Barrio-Hernandez, I. et al. Clustering predicted structures at the scale of the known protein universe. Nature 622, 637–645 (2023).

24. Farrar, M. Striped Smith–Waterman speeds database searches six times over other SIMD implementations. Bioinformatics 23, 156–161 (2007).

25. Remmert, M., Biegert, A., Hauser, A. & Söding, J. HHblits: lightning-fast iterative protein sequence searching by HMM-HMM alignment. Nature Methods 9, 173–175 (2012).

26. Steinegger, M. & Söding, J. Clustering huge protein sequence sets in linear time. Nature Communications 9, 2542 (2018).

27. Mariani, V., Biasini, M., Barbato, A. & Schwede, T. lDDT: a local superposition-free score for comparing protein structures and models using distance difference tests. Bioinformatics 29, 2722–2728 (2013).

28. Armougom, F., Moretti, S., Keduas, V. & Notredame, C. The iRMSD: a local measure of sequence alignment accuracy using structural information. Bioinformatics 22, e35–e39 (2006).

29. Mizuguchi, K., Deane, C. M., Blundell, T. L. & Overington, J. P. HOMSTRAD: A database of protein structure alignments for homologous families. Protein Science 7, 2469–2471 (1998).

30. Lau, A. M. et al. Exploring structural diversity across the protein universe with the encyclopedia of domains. bioRxiv 2024.03.18.585509 (2024).

31. Orengo, C. et al. CATH – a hierarchic classification of protein domain structures. Structure 5, 1093–1109 (1997).

32. Puente-Lelievre, C. et al. Tertiary-interaction characters enable fast, model-based structural phylogenetics beyond the twilight zone. bioRxiv 2023.12.12.571181 (2024).

33. Chin, D. & Means, A. R. Calmodulin: a prototypical calcium sensor. Trends in Cell Biology 10, 322–328 (2000).

34. Tidow, H. & Nissen, P. Structural diversity of calmodulin binding to its target sites. The FEBS Journal 280, 5551–5565 (2013).

35. Li, Z., Jaroszewski, L., Iyer, M., Sedova, M. & Godzik, A. FATCAT 2.0: towards a better understanding of the structural diversity of proteins. Nucleic Acids Research 48, W60–W64 (2020).

36. Wilson, M. T. & Reeder, B. J. in MYOGLOBIN (eds Laurent, G.J. & Shapiro, S.D.) Encyclopedia of Respiratory Medicine 73–76 (Academic Press, Oxford, 2006).

37. Moi, D. et al. Structural phylogenetics unravels the evolutionary diversification of communication systems in gram-positive bacteria and their viruses. bioRxiv 2023.09.19.558401 (2023).

38. Mifsud, J. C. O. et al. Mapping glycoprotein structure reveals defining events in the evolution of the Flaviviridae. bioRxiv 2024.02.06.579159 (2024).

39. Heinzinger, M. et al. Bilingual Language Model for Protein Sequence and Structure. bioRxiv 2023.07.23.550085 (2024).

40. Kim, W. et al. Rapid and Sensitive Protein Complex Alignment with Foldseek-Multimer. bioRxiv 2024.04.14.589414 (2024).

## References

41. Gotoh, O. An improved algorithm for matching biological sequences. Journal of Molecular Biology 162, 705–708 (1982).

42. Gotoh, O. Significant improvement in accuracy of multiple protein sequence alignments by iterative refinement as assessed by reference to structural alignments. Journal of Molecular Biology 264, 823–838 (1996).

43. Mirdita, M., Steinegger, M. & Söding, J. MMseqs2 desktop and local web server app for fast, interactive sequence searches. Bioinformatics 35, 2856–2858 (2019).

44. Stebbings, L. A. & Mizuguchi, K. HOMSTRAD: recent developments of the Homologous Protein Structure Alignment Database. Nucleic Acids Research 32, D203–D207 (2004).

45. Notredame, C., Higgins, D. G. & Heringa, J. T-coffee: a novel method for fast and accurate multiple sequence alignment. Journal of Molecular Biology 302, 205–217 (2000).

46. Lau, A. M. et al. Exploring structural diversity across the protein universe with the encyclopedia of domains. bioRxiv 2024.03.18.585509 (2024).

47. Orengo, C. et al. CATH – a hierarchic classification of protein domain structures. Structure 5, 1093–1109 (1997).

48. Berman, H. M. et al. The Protein Data Bank. Nucleic Acids Research 28, 235–242 (2000).

49. Meng, E. C. et al. UCSF ChimeraX: Tools for structure building and analysis. Protein Science 32, e4792 (2023).

50. Mifsud, J. C. et al. Underlying data for “mapping glycoprotein structure reveals flaviviridae evolutionary history” (2024). URL https://zenodo.org/records/11092288.

51. Mifsud, J. C. O. et al. Mapping glycoprotein structure reveals defining events in the evolution of the Flaviviridae. bioRxiv 2024.02.06.579159 (2024).

52. Capella-Gutierrez, S., Silla-Martinez, J. M. & Gabaldon, T. trimal: a tool for automated alignment trimming in large-scale phylogenetic analyses. Bioinformatics 25, 1972–1973 (2009).

53. Kalyaanamoorthy, S., Minh, B. Q., Wong, T. K. F., von Haeseler, A. & Jermiin, L. S. Modelfinder: fast model selection for accurate phylogenetic estimates. Nature Methods 14, 587–589 (2017).

54. Minh, B. Q. et al. IQ-TREE 2: New Models and Efficient Methods for Phylogenetic Inference in the Genomic Era. Molecular Biology and Evolution 37, 1530–1534 (2020).

55. Hoang, D. T., Chernomor, O., von Haeseler, A., Minh, B. Q. & Vinh, L. S. UFBoot2: Improving the Ultrafast Bootstrap Approximation. Molecular Biology and Evolution 35, 518–522 (2018).

56. Xu, S. et al. Ggtree: A serialized data object for visualization of a phylogenetic tree and annotation data. iMeta 1, e56 (2022).

57. Sand, A. et al. tqDist: a library for computing the quartet and triplet distances between binary or general trees. Bioinformatics 30, 2079–2080 (2014).

58. Smith, M. R. Quartet: comparison of phylogenetic trees using quartet and split measures (2019). R package version 1.2.6.9001.

